# “The best home for this paper”: A qualitative study of how authors select where to submit manuscripts

**DOI:** 10.1101/2024.05.14.594165

**Authors:** Lauren A. Maggio, Natascha Chtena, Juan Pablo Alperin, Laura Moorhead, John M. Willinsky

**Author notes:** Correspondence should be addressed to Lauren A. Maggio, Department of Medical Education, University of Illinois at Chicago, 808 S Wood St., Chicago IL 60612;. Funding Support*: Anonymous donor, Stanford University. Ethical Approval:* Stanford University IRB (Protocol#70447) and Simon Fraser REB (Case #30001784). Disclosures:* None reported. Data:* None reported.

## Abstract

**Introduction:** For academics selecting a target journal to submit a manuscript is a critical decision with career implications. In medical education, research conducted in 2016 found that authors were influenced by multiple factors such as a journal’s prestige and its mission. However, since this research was conducted the publishing landscape has shifted to include a broader variety of journals, an increased threat of predatory journals, and new publishing models. This study updates and expands upon how medical education authors decide which journal to submit to with the aim of describing the motivational factors and journal characteristics that guide authors’ decision making.

**Methods:** The authors conducted five qualitative focus groups in which twenty-two medical education authors and editors participated. During the focus groups participants were engaged in a discussion about how they select a journal to submit their manuscripts. Audio from all focus groups was transcribed. Transcripts were analyzed using codebook thematic analysis.

**Results:** Participants considered multiple factors when selecting a target journal. Factors included a journal’s impact, the scope of a journal, journal quality, and technical factors (e.g., word limits). Participants also described how social factors influenced their process and that open access plays a role that could both encourage or deter submission.

**Discussion:** The findings describe the motivational factors and influential signals that guide authors in their journal selection decision making. These findings confirm, extend, and update journal selection factors reported in medical education and other disciplines. Notably, these findings emphasize the role of social factors, relationships and personal experiences, which were absent from previous work. Additionally, we observed increased consideration of OA and a shift away from an emphasis on journal prestige.

---

Where researchers decide to submit their manuscripts has significant consequences for their careers. Choosing a journal is not an easy choice, when considering the common estimate of there being over 33,000 academic journals available this is not a simple choice^1^ or even more so if you are persuaded by those who regard this figure as a vast and prejudicial undercount^2^. Choices only become more challenging as the publishing landscape continues to change with the “publish or perish” culture^3^, the growth of open access^4^, and the rise of publishing models like preprints and post-publication review^5^. Medical education publishing has not been immune to these changes with the introduction of new journals and publication models^6-9^ and the transition of some journals from a subscription model to open access^10^.

The journal to which an academic submits their scholarship influences their career trajectory. For example, selecting one journal over another may result in a lengthier publication timeline than had another been chosen^11^ or could mean whether or not an article counts for an author’s promotion^12^. This is underscored by the fact that the majority of medical school promotion guidelines endorse publishing in “prestigious” or “high-impact” journals and stress publication quantity^12^. Complicating matters, this emphasis on prestige and publication quantity has been linked to publication pressure, which can open the door to so-called “predatory” journals. Predatory journals “prioritize self-interest at the expense of scholarship and are characterized by false or misleading information, deviation from best editorial and publication practices, a lack of transparency, and/or the use of aggressive and indiscriminate solicitation practices”^13 p.211^. Researchers have found that physicians and medical trainees struggle to identify predatory journals, which can damage researchers’ credibility and waste resources^14,15^. These high stakes suggest that a better understanding of how authors select journals is warranted.

In 2022, Olle ten Cate, a senior medical education scholar, published a commentary in which he answered the frequently asked question, “Can you recommend a journal for my paper?,”^16^ suggesting that medical educators are interested in further guidance on this important topic. ten Cate’s response draws on personal experience and a recent study that surveyed medical educators about journal choice^17^. Choosing from fixed responses, study respondents selected fit, impact, and editorial reputation as the top drivers of their decisions. While valuable and more current, and in some cases aligned with earlier qualitative research by Ginsburg et al.^18^, this survey study provides limited insight into how authors consider the factors that motivate their decision to target a particular journal. Thus, in light of the importance of journal selection and the evolving landscape, this study uses author and editor perspectives to revisit how medical education authors decide which journal to submit to with the aim of providing the motivational factors and influential signals that guide their decision making. Our aim is that these findings will facilitate new and seasoned researchers in navigating the evolving complexities of journal selection and also alert editors to those factors and signals, such that they can be highlighted in their journal processes.

## Methods

We conducted qualitative focus groups to understand how HPE authors and editors select journals to submit articles for publication. Our study design and interpretation of the data are guided by a constructivist paradigm, through which knowledge is perceived as socially constructed between an individual and society^19^. Stanford University (Protocol#70447) and Simon Fraser University (Case#30001784) granted ethical approval to conduct this study.

### Data Collection

Our study focused on authors and editors who had published in or oversaw medical education journals. We defined medical education journals as those included on the Medical Education Journals List (MEJ-24)^20^.

To identify authors, we searched PubMed for articles published in the MEJ-24 in 2022. Then using a random-number generator, we selected 50 articles and extracted the first author’s name and email. Authors without listed email addresses were excluded. Over the recruitment period, we continued this process until we had recruited enough participants to reach information power based on our research aim^21,22^. We focused on first authors publishing in 2022, as they had recently navigated and likely led the journal selection process.

For the editors, we included the Editor-in-Chief (EIC) for MEJ-24 journals and their deputy editor(s). All editor participants had authored at least one article in an MEJ-24 journal. One journal, *BMJ Simulation and Technology Enhanced Learning*, ceased publication in 2023. We excluded this journal’s editors, but included its authors. Between June and November 2023, LAM, a medical education author and Deputy EIC of *Perspectives on Medical Education*, emailed editors to participate.

Data was collected using focus groups. Focus groups can engage participants in conversations that explore their understanding of a topic, as well as their views about that topic^23^. We conducted five focus groups: one included only editors and four included a combination of authors (who were not editors) and editors. Researchers have found that homogeneity of participants within focus groups can provide a safe environment that mitigates a sense of power imbalance^24^. As such, we initially separated authors and editors into different groups to determine whether a difference in discussion existed. However, we did not observe a difference and conducted the remaining focus groups with a combination of participants.

LAM and NC, a researcher with experience in qualitative research and a background in education, information studies and optometry, moderated the focus groups using a semi-structured guide (see Appendix 1). We piloted the guide with a group of six medical educators, including an editor, and minor modifications were made based on participant feedback. In the focus groups, LAM and NC posed open-ended questions that invited participants to reflect on their journal selection process. Focus groups were 75 minutes on GoogleMeets, with participants logging in from different locations. All focus groups were audio recorded and transcribed using GoogleMeets. LAM and NC checked the transcripts for accuracy.

### Analysis

Transcripts were analyzed using codebook thematic analysis^25^. LAM and NC independently coded all five transcripts initially using the themes identified in Ginsburg’s interview study conducted in 2016^18^ as predetermined codes to guide analysis, while remaining open to additional codes. After initial coding, they met several times to review and discuss the code book with revisions made as necessary. Upon agreement, LAM and NC applied the codes to all transcripts and presented the author team with identified themes and representative quotes for review and refinement until consensus was achieved.

### Reflexivity

LAM is a medical education researcher and editor. LAM knew many of the participants as colleagues, which may have influenced results, as respondents may have been inclined to provide responses pleasing to her, or may have withheld information for fear of being judged by “one of their own.” This was reflected upon both during the interview process and during data analysis and interpretation. Fellow author team members with expertise in education and scholarly publishing, but who were unfamiliar with medical education and the participants, provided a counterbalance to potential bias, which was taken into consideration in the analysis and reporting of results.

## Results

Twenty-two individuals participated with focus groups ranging in size from 4–5 participants. Participants primarily were professors at the ranks of assistant (n=3), associate (n=4), full (n=11), but also included three PhD candidates and an academic librarian. Participants were based in the United States (n=6), Canada (n=5), Australia (n=3), Netherlands (n=2), and one participant each from Austria, Barbados, Norway, and Singapore. Nine editors participated.

Multiple factors influenced participants’ journal selection. Overall, factors were similar across the five focus groups, including the group composed solely of editors. However, we did appreciate differences based on participant seniority, which we note in relation to specific factors.

### Impact

Participants strongly considered a journal’s *impact* when selecting a journal. Generally, participants described impact as a journal’s ability to reach a broad and desired readership. “When I publish in *Academic Medicine*, I know the right people are really reading that journal, so it will have impact” (5A). Another participant described transitioning a qualitative research article to a non-research publication to satisfy the editors of a “highly influential” journal:

> Basically, we submitted it as a piece of original research and they wrote back and said, “You’ve called it research, but it’s qualitative.” And we should have said this is terrible. Lots of red flags, but this is a hugely influential journal…in some ways we just held our noses and said, this is going to reach a wide readership. (E2)

Participants also discussed impact in relation to the journal impact factor (JIF), which is a citation-based metric. The role of the JIF in journal selection varied based on career stage, institutional context, and geographical setting. Overall, senior scholars were more likely to report not considering JIF as a motivation in their decision-making, though one PhD student and the librarian also mentioned ignoring a journal’s JIF. One full professor noted, “I’m at the stage in my career where I really don’t care about impact factors [laughing]. I’m actually more about getting something read” (1D). Participants who ranked the JIF low in their selection criteria—or who reported not including it at all—made comments about reaching the right readership as outweighing prestige: “[In] my department, we don’t care as much about impact factor or things like that, as much as we care about is it going to get to the right audience” (4A).

However, not everyone downplayed the JIF. A minority of participants acknowledged that publishing in journals with a high JIF improved their work’s reach and impact: “I’ve personally never felt any pressure to publish in a journal because of impact factor…But at the same time, I do feel a little bit of interest in publishing in high impact journals. The only reason is that I know that it’s read” (5A).

### Fit

Participants seriously considered whether their article was in a given journal’s scope. This consideration was tightly linked to the perceived likelihood of an article being accepted in that journal. Participants mentioned looking at mission statements and objectives to identify “what is going to be attractive to a journal” (2D) and whether “there would be a welcoming audience” (E5). Relatedly, participants mentioned reading the author guidelines, as well as the journal’s content to “see what they’re publishing”(1D) in order to gauge suitability and fit. Participants also used these strategies to identify a journal’s audience. One participant explained, “it’s definitely about who seems to be the dominant readership of that journal. Are they the people that are going to be interested in the conversation I’m trying to advance?” (E3).

Participants also considered the match with their study’s methodology, context and goals when weighing journal options:

> So I would think: Where did we do this study? Was it in medicine? Was it post-graduate, undergraduate? So, then, I would think is it a very practical type of study? Or is it really a study or is it guidelines or something? […] And then you could kind of weigh in, what would be journals that you would expect would more easily accept this paper because at the end, you want most papers to really have a home. (5A)

Interestingly, while participants considered their study’s perceived quality and rigor in the selection process, the way that it impacted journal choice was not always straightforward. In discussing how journal choice is negotiated within her author team, one participant noted, that clinical journals were sometimes perceived as more welcoming than medical education journals:

> I think it’s interesting that sometimes the higher impact journals that people might be aiming for are actually not medical or health professions education journals. They are the clinical journals in their field…They might actually be more inclined to publish really pragmatic pieces of work about, “We did this thing, and you can do it too.” Stuff that some of our more familiar journals would say, “That doesn’t really meet our bar.” …And so, we often talk about, “Who are you hoping to influence here?” If my co-authors want to influence pediatricians, who are trying to teach professionalism, then I’m happy for them to try to put it in a pediatrics journal. That wouldn’t be probably where I would put my work primarily, but it might be the audience they want to talk to. (E2)

Participants considered geographic context often seeking to match studies conducted in a particular country or region with a journal’s scope.

### Journal Technical Factors

Participants mentioned multiple journal technical factors, including the journal’s publication timeline, word limits, and where it is indexed. Time to publication was a major factor for many participants. On one hand, a participant noted, “If I know a journal probably will get back to me quickly and it does fit, then maybe I’ll try that even if I think it might be a bit of a reach.” (1B) On the other hand, an editor said:

> There’s nothing more damaging to a journal than the interpretation of being very, very slow to get a decision back. And so there are certainly journals I avoid. I avoid them purely because there’s every indication from people in the communities that it’s taking a year to get things turned around and that’s not good for anyone. (E3)

Publication timelines were especially “crucial” (2C) for doctoral students on a graduation timeline. Participants acknowledged that journals had elongated publication timelines for a variety of reasons outside editorial control (e.g., difficulties securing peer reviewers, slow resubmission revisions).

Participants discussed selecting journals based on word limits with an aim of avoiding those they considered too restrictive. One participant described a journal: “It’s got a 1500 word cap, and I don’t know what you can say, in 1500 words that it doesn’t come across like a haiku.” (E4) Some participants also mentioned a journal’s indexation (e.g., included in MEDLINE), which was important for an article’s findability, but also seen as a reflection of a journal’s quality and important for promotion.

### Peer Review

Participants factored the quality and process of peer review into submission decisions with “knowing that it [a manuscript] gets a real peer review by experts” (4C) being critical.

Participants’ perceptions of a journal’s peer review processes, which were likened to a quality assurance process, were shaped by their experiences as peer reviewers and authors.

> I’ve had a couple [of experiences] where the papers that were sent to me were ones that I actually think shouldn’t have even made it to review. They were of such poor quality. And then another instance is when you get back the copy of what all the reviewers said. Sometimes my review will be the most detailed one there. There’s sometimes others that have written a paragraph and they don’t seem like an expert in the field at all, or anyone who has any knowledge of health professions education. So those would be red flags to me as a reviewer or I guess if I was an author and I got back reviewer comments like that, that would kind of be a flag. (1A)

As peer reviewers, participants were attuned to the editor’s role in thoughtfully assigning peer reviewers (e.g., not sending a quantitative researcher a qualitative paper) and helping authors make sense of multiple reviews by presenting a synthesis in the decision letter. When the participants felt the editor acted poorly, it negatively impacted their decision to submit future manuscripts to the journal.

In preparation to submit, participants described reading potential target journals and noting that certain characteristics made them question the quality of peer review. For example, participants described that for those journals that published their article’s publication timeline that “a really, really short time between acceptance and between submission” (E1) suggested poor quality peer review. Participants also mentioned that if a journal published “a mixed bag quality of papers” (4D) that they would question its peer reviewer process and likely avoid submitting to that journal in the future.

### Open Access

Participants’ attitudes towards open access (OA) as a journal selection factor varied depending on the journal’s perceived quality and reputation, the journal’s editorial and peer review practices, and the cost of article processing charges (APCs). Overall, participants described OA in terms of its benefits, including increased citation counts, copyright retention, and ability to reach scholars with limited access (e.g., Global South researchers). OA was generally not a driving factor in decision-making, but a complementary one:

> “I’m not sure that I’ve chosen a journal because my institution’s relationship allows open access, but I do see it as a plus. Okay, I wouldn’t say that it was a decision-maker in terms of where a paper ended up going, but perhaps it just strengthened that decision.” (1B)

Participants with institutional support for APCs or large research budgets viewed OA more favorably and factored it into their decision-making process, as did those with a strong belief in OA’s emancipatory effects. One participant’s experience as a researcher in the Global South inclined them to view OA as a way to expand their work’s reach and to bridge knowledge divides: “Sometimes I think about open access as my first choice, maybe because I’m coming from a low-resource country. And I know how people there suffer in finding the information […] Sometimes I think, if I can publish in an open access journal, then people can have it.” (3B)

At the same time, several participants mentioned needing to balance publishing OA with APCs. For those without institutional OA budgets, APCs were a major deterrent:

> Open access fees can absolutely be a barrier […] I simply won’t consider those journals because I just can’t [pay] those fees so often, even if it’s probably the best journal for me to submit an article to. I absolutely just can’t. And that’s a huge barrier in particular for graduate students. (2C)

Additionally, one participant mentioned how excessive APCs can raise questions about a journal’s integrity: “When I see the APCSs that are very high I consider: Are they running this journal just to earn money or is it a reliable journal?” (1C). Some participants also linked high APCs to predatory publishing. Lastly, a few participants noted how colleague’s outdated notions about OA can undermine work published in legitimate OA journals because these are wrongfully perceived as predatory or low quality:

> They will say, “Ohh, you’re paying to publish,” as if it’s something [you] shouldn’t be doing because they don’t understand what’s been happening in publishing over the last five to 10 years. So, you know, you do have to navigate through this. (5A)

### Social Factors / Relationships

Several participants highlighted the role of professional relationships and experience. Multiple senior authors, defined as those at the professor level, mentioned knowing many journal editors personally and being able to “just approach them” (5A) to inquire if a particular manuscript would be of interest. They also discussed how journal selection is guided by their maturity of experience and familiarity with a journal more broadly: “When you’ve been around for long enough, you know who is reading them to some extent, you know who the editors are.” (4D)

On this theme, a participant described that they keep the field’s leading editors in mind as they execute studies or write up findings. Notably they referred to these editors on a first-name basis, indicating familiarity and, arguably, a privilege of proximity that most authors do not have: “I have this voice in my head when I’m writing things or doing studies. And I think Kevin will like this or Rachel will like this or Erik might like this. Or, equally, I might have in my head: ‘No, there’s no point. Kevin won’t like this.’” (2D)

On the flipside, some junior and mid-level scholars mentioned their lack of connections and insider knowledge. One participant reflected on her struggles as a novice author after hearing the more senior authors in the group discuss their relationships with editors and how these have informed their journal publishing experience:

> It’s been a very interesting conversation to listen to such esteemed medical educators and researchers, because as somebody who transitioned into this field, it did feel challenging to get through the barrier of what I perceived as a network that you couldn’t break through […] It feels like if you’re not in the know, you’re going to be marginalized. (3D)

Furthermore, participants noted that personal experience as an editor and/or peer reviewer factored into the journal selection process. One participant mentioned that as “an editor, you do get a lot of insider information” (2D), while another said: “I’ve been a reviewer for a lot of journals. So I know, sort of the quality of reviews and the timelines just based on personal experience.” (1A) Several participants shared the sentiment that being an editor and peer reviewer offered “intelligence” not readily available otherwise. One participant mentioned how serving as peer reviewer can be the “start of a relationship” (3C) with a journal.

Finally, some participants mentioned mentorship in the context of journal selection, either as providers or receivers. One participant noted the criticality of mentorship, adding that without a mentor they “wouldn’t have known where to go” (3D) to publish.

## Discussion

Our findings describe the motivational factors and influential signals that guide authors in their decision making. These findings confirm, extend, and update journal selection factors reported in medical education^17,18^ and beyond^26,27^. In medical education, our findings reinforce previously identified factors such as fit with the scope of the journal, journal quality and reputation, journal audience, and technical factors (e.g., turnaround time). However, there are also deviations.

Notably, our findings emphasize the role of social factors, relationships and personal experiences in journal selection. Additionally, we observed increased consideration of OA and a shift away from an emphasis on journal prestige (ie., JIF).

The role of social factors and relationships in journal selection were absent from earlier medical education studies^17,18^ and only mentioned peripherally in other disciplines and contexts. For example, an interdisciplinary survey of international academics found that having a relationship with a journal influenced journal choice, but relationships were described more formally such as serving on the journal’s editorial board^27^. These relationships feel distinct from the personal relationships and first-name familiarity mentioned by our senior participants, both editors and non-editors alike. This finding is important because it suggests that less experienced, less in-the-know scholars with smaller and/or weaker networks may be significantly disadvantaged in the publishing process. While perhaps not surprising, this finding stands out in a professional and educational landscape increasingly tasked with improving diversity and addressing issues of equity in medical education publishing^28,29^. This finding signals that authors, especially those new to the field, may benefit from increased opportunities to engage with and build relationships with journals and editors. Journals could facilitate these relationships by being explicit about how individuals can become involved with the publication. For example, journal websites could prominently feature opportunities to become a peer reviewer and advertise calls for editorial board positions with clear criteria for such roles. Additionally, journals might consider adopting and building upon existing initiatives. For example, the journal *Medical Education* recently instituted “online office hours” which encourage authors to directly engage with its editors in an informal setting and are held across multiple timezones to accommodate a global audience ^30^. Another example is *Academic Medicine*’s creation of a path towards editorship program, which welcomes and trains early and mid-career scholars^31,32^.

All focus groups mentioned OA as a consideration, but not a driving force in journal selection, which aligns with studies across disciplines^26,27,33^. However, unlike in Ginsburg et al.’s^18^ medical education study in which participants expressed apprehension toward OA many of our participants looked favorably on it, citing the increased reach and global unfettered exposure it provides. Negative comments overwhelmingly focused on affordability—specifically, the high cost of APCs, which participants noted disproportionately affect ECRs and those who lack access to funds for APCs. Overall, our findings suggest that OA publishing has, in recent years, gained traction in medical education—likely due to institutions engaging in transformative agreements directly with publishers, funder mandates, and the impact of events like COVID-19 on scientific practice and communication^34^. However, there are still misconceptions about OA and financial barriers to be addressed. For misconceptions about OA, many resources are available to educate researchers, including those specific to medical education (e.g., Open Access: What it Means for Your Article^35^). APCs are indeed a barrier, especially in a field with increasing, but still limited funding^36^. To shift the burden from authors, some institutions have engaged in agreements with publishers that bundle subscriptions with the APCs or discounted APCs for the institution’s authors. For example, at the University of Chicago, an institutional agreement allows authors to publish OA without facing an APC in Wiley journals, which includes several relevant titles (e.g., *Medical Education, Academic Emergency Medicine)*^37^.

In Ginsburg et al.’s study^18^, JIF was identified as the most prevalent factor in journal selection and was considered an important priority for those in Rees’ et al.s survey^17^. However, in our findings JIF was deemed less important though still acknowledged as influential in relation to readership. Related to JIF, institutional policies regarding the ranking of journals and articles were viewed as relatively important, but not as important as, for example, the quality, integrity, and reliability of the reviewing process. It is possible that this is the fruit of global initiatives to reform academic assessment and deemphasize the JIF, such as the Declaration on Research Assessment^38^. Our finding may reflect that many of our participants were senior scholars, which also aligns with Niles et al.’s finding that the JIF was less important to older and likely more senior participants^26^. Future research might further investigate the perceptions of early career researchers to better understand their stance on JIF and other related metrics.

### Limitations

Our findings are context specific, which may limit their generalizability outside their context. While we included scholars from 8 countries, representation was lacking from certain regions (e.g., South America). Future researchers might consider a more targeted approach to recruiting a broader diversity of authors. Additionally, recruitment focused on individuals that had published in MEJ-24 journals, which is an incomplete representation of medical education (Maggio, 2023). However, our participant population roughly aligns with studies describing medical education authors^39,40^. Also, our inclusion of journal editors may have skewed our study population towards more advanced researchers. Future researchers could investigate this topic with other populations.

## Conclusion

This study provides an international perspective on the factors and signals that medical education researchers consider when selecting a journal for their research. Participants in the study demonstrated a thoughtful and nuanced approach considering various journal’s impact, audience, and accessibility, and made decisions based on the alignment between the journal and their research goals and values.

## Appendix Semi-structured focus group guide

### To get started, imagine that you are preparing to submit a manuscript focused on an HPE topic. Please describe how you would select your target journal

Potential Prompts (not all questions will necessarily be asked)

- What specific factors come into play and why are they important?
  ∘ *Likely journal impact factor (JIF) will be mentioned. If necessary, after a few minutes of discussion about the JIF state: Taking impact factor off the table, which other factors are important to you?*
  ∘ How does your institution’s promotion and tenure guidelines influence your decision?
  ∘ In previous studies, HPE authors have said that access to their work is important. What role, if any, does a journal’s open access status play?
    ▪ What about if there is an author processing charge?
- Before submitting to a journal, what are some of the factors that you wished you knew about that are not readily accessible? *If an example is needed, you could say rejection rate*.
- What roles do your co-authors play in this decision?
  ∘ At what point do you discuss your target journal with your team?
  ∘ In what ways, if any, does the makeup of your team influence selection (e.g., having ECRs, someone going up for tenure/promotion, clinically based authors)
- When do you consider publishing in a specialty academic journal (e.g., Academic Pediatrics) vs. a more general HPE journal?
  ∘ In what ways, if any, would your approach be different if you were planning to publish outside of HPE, for example in a clinical or psychology journal?
- How does your process change after a rejection? After the third rejection?
- Previous research has shown that a journal’s “reputation” plays into journal selection decisions. Do you feel this is still true and if yes, how do you judge a journal’s reputation?
  ∘ What do you consider a red flag and would it stop you from submitting?
  ∘ What are characteristics of a journal that reassure you about its integrity?
  ∘ Where do you look for information about a journal’s integrity?
  ∘ In what ways, if any, do you guard against submitting to predatory journals?
  ∘ How did you learn about journal integrity?

